# A method for evaluating hunger and thirst in monkeys by measuring blood ghrelin and osmolality levels

**DOI:** 10.1101/2023.11.06.565780

**Authors:** Yuki Suwa, Jun Kunimatsu, Akua Kamata, Masayuki Matsumoto, Hiroshi Yamada

## Abstract

Hunger and thirst drive animals’ consumption behavior and regulate their decision-making regarding rewards. We previously assessed the thirst states of monkeys by measuring blood osmolality under controlled water access and related the thirst states to risky decisions for fluid rewards. However, hunger assessments in monkeys have been poorly performed. Moreover, the lack of precise measures for hunger states leads to another issue regarding how hunger and thirst states interact with each other in each individual. Thus, when controlling food access to motivate subject performances, how these two physiological needs are satisfied in captive monkeys remains unclear. Here, we measured blood ghrelin levels and osmolality for hunger and thirst, respectively, in four captive macaques. Using enzyme-linked immunosorbent assay, we found that the levels of blood ghrelin, a widely measured hunger-related peptide hormone in humans, were high after 20 h of no food access (with *ad libitum* water), which is a typical controlled food access condition. One hour after consuming the regular dry meal, the value decreased in three out of four monkeys to within the range of individual blood ghrelin levels. Additionally, blood osmolality measured from the same blood sample, the standard hematological index of hydration status, increased after consuming regular dry meals with no water access. Thus, ghrelin and osmolality may provide a precise reflection of the physiological states of individual monkeys for hunger and thirst, suggesting that these indices can be used as a tool for monitoring hunger and thirst levels that mediate the animal’s decision-making for consuming rewards.

## Introduction

Hunger and thirst are fundamental constituents of physiological needs that drive an animal’s consumption behavior to maintain its physical state (Mattes, 2010). Controlled access to food or beverages is utilized in standard experimental procedures for non-human primates, which motivates subject performance. Hunger drives the consumption of food that maintains the metabolic state, and thirst drives the intake of fluid that maintains the hydration state (Berne and Levy, 1993), although these physiological needs are not independent with each other (e.g., food items contain both energy and fluid) (Ramachandran and Pearce, 1987; Watts, 1999; Betley et al., 2015). For example, thirsty animals are unlikely to eat dry food, even if they are hungry (Eiselt et al., 2021), which implies that the reward values of food and beverages may be somehow linked in the reward circuitry (Haber and Knutson, 2010). Hunger and thirst levels are subjectively evaluated and controlled by experimenters, although these cravings in each individual differentially change owing to variability in the individual homeostatic state. Thus, establishing a method to reliably evaluate the hunger and thirst states of animals is useful, which drives their consumption behavior to appropriately maintain their physical state.

In the experimental testing of behavior in monkeys and rodents, a typical procedure controls either of these physiological needs to motivate the animal’s performance, especially in experiments involving electrophysiology in monkeys (Evarts, 1968; Wurtz, 1968; Watanabe, 1996). To assess thirst, the blood osmolality level can be used to detect the level of hydration status in mammals as the standard hematological index (Rolls and Rolls, 1982; Stedman, 2006). We have previously used the osmolality level to relate the behavioral characteristics of economic decisions, such as risk preferences (Yamada et al., 2013), instrumental performance of actions to earn fluid rewards (Minamimoto et al., 2012), and simple eye movements for fluid reward intake (Yamada et al., 2010) in monkeys, a close relative of humans. To assess hunger, various physical measures are used (Mattes, 2010), such as blood sugar levels and insulin as standard measures and leptin and ghrelin as recently developed measures. These physical indices of hunger and thirst are useful for assessing and controlling the motivational state of subjects. However, the lack of a simultaneous assessment of hydration and metabolic status leads to difficulty in evaluating neural correlates for hunger and thirst in behaving monkeys because they are related and control of either one mostly affects the other status (Ramachandran and Pearce, 1987; Watts, 1999; Betley et al., 2015).

This limitation induces several potential problems in Neuroeconomic studies (Camerer et al., 2005; Glimcher et al., 2008), which employ experimental testing of reward valuation systems for economic choices. When measuring neural activity in reward circuitry, the subjective values of any reward depend on the physical state of the subject (Nakano et al., 1984; Critchley and Rolls, 1996; de Araujo et al., 2006; Pritchard et al., 2008), even for money (Symmonds et al., 2010). Because hunger and thirst may dramatically change neural activity in most parts of the brain (de Araujo et al., 2006), the control and monitoring of either need is insufficient for precisely assessing the neural valuation system embedded in the reward circuitry. The subjective values of the items used in the monkey experiments depend on these physiological statuses; for example, the intake of juice rewards elicits changes in both statuses. Thus, assessing both hunger and thirst states under controlled access to either food or beverages is worthwhile.

Herein, we measured blood ghrelin levels for assessing hunger (Kojima et al., 1999) and osmolality for assessing thirst (Rolls and Rolls, 1982) in captive monkeys. Using a typical experimental procedure to control food and fluid access, we evaluated the physical status of monkeys. Our results indicated that these indices can be used to monitor hunger and thirst levels in individual monkeys.

## Materials and Methods

### Experimental subjects

Four macaque monkeys were used in this study (*Macaque fuscata*; monkey MON, male, 7.8 kg, 4 years; monkey Y1, male, 6.4 kg, 2 years; monkey MIY, female, 5.8 kg, 4 years; *Macaque mulatta*; monkey SUN, male, 7.5 kg, 13 years). The Animal Care and Use Committee of the University of Tsukuba approved all experimental procedures (protocol no. H30.336), which were also performed in compliance with the US Public Health Service’s Guide for the Care and Use of Laboratory Animals.

### Blood Collection

The monkeys were anesthetized using medetomidine (0.03 mg/kg, intramuscularly [i.m.]) and midazolam (0.3 mg/kg, i.m.), except monkey SUN who was seated in a standard primate chair and desensitized to leg restraint by using positive reinforcement with food rewards before starting the blood collection procedure. Blood samples (2.0 ml) were drawn from the saphenous vein using a butterfly needle (22 gauge) with a single collection. Atipamezole (0.024 mg/kg, i.m.) was administered and 30 min after recovery of the monkeys, they were fed a dry meal. Approximately 2 h after the first blood collection, the remaining food was extracted. After 1 h (a total of 3 h spent for food intake and the following rest in the home cage), a second blood sample was corrected using the same procedure as for the first sample collection. The monkey SUN was sampled while awake, and after desensitization, exhibited no distress during the sampling procedure. Approximately 1.0 ml plasma was extracted in each collection from the 2.0 ml blood sample in ethylenediaminetetraacetic acid-2K-containing tubes by centrifugation at 2000 × *g* for 10 min at 4°C. The total amount of blood extracted within any two-week experimental period did not exceed 5% of the total blood volume (total blood volume was estimated at 65 ml/kg weight).

A total of 16 blood collections were performed in eight days for each monkey (two blood collections per day before and after food intake). A total of 64 blood samples were collected from the four monkeys.

### Controlled access to food and water

In the approved controlled food access protocol, the monkeys received a fixed daily allocation of dry meals (certified diet, PS-A, Oriental, Japan). This daily food was allocated at least once a day at approximately noon, except for the periods before and after surgery. The fixed daily allocation of food was as follows: monkey MON, 160 g or 21 g/kg/day; monkey Y1, 140 g or 22 g/kg/day; monkey MIY, 130 g or 22 g/kg/day; and monkey SUN, 140 g or 19 g/kg/day. Under the experimental conditions employed, we removed all leftover food at 14:00 on the day before blood collection, which led to approximately 20 h of no food access. Food was subsequently delivered after the first blood collection, with the amount allocated daily to each monkey at approximately 10:00. The average amount of food consumed by each monkey during the test period was as follows: monkey MON, 159 g or 20 g/kg/day; monkey Y1, 140 g or 22 g/kg/day; monkey MIY, 127 g or 22 g/kg/day; and monkey SUN, 134 g or 18 g/kg/day. If the monkey did not quickly start eating the dry meal, small pieces of sweet potatoes were fed to the monkey (< 10 g). The monkeys were free to access a 500-ml bottle of water in a day, and no water access control was performed throughout the entire period of the test that the monkeys engaged in, except for the period between the first and second blood collections without water access. This procedure mimics a typical experimental procedure to control food access that motivate the subject performance to obtain food rewards in the experimental room.

### Body weight measurements

The body weight of each monkey was measured before the first blood collection.

### Measuring the blood ghrelin levels using enzyme-linked immunosorbent assay (ELISA)

Unacylated ghrelin levels were measured using ELISA (Unacylated Ghrelin (human) Express Enzyme Immunoassay kit, Bertin Pharma, #A05119.96 wells, France). The immunoassays were performed according to the manufacturer’s instructions. The plates coated with the primary antibody in each well were rinsed five times with the wash buffer (300 µL/well) included in the kit. The standards, controls, and samples were added to the plates. Additionally, 100 µL of a tracer was dispensed to the wells, and the plates were incubated at room temperature for 3 h (RT, approximately 20 ∼ 25 °C). After rewashing, 200 µL of the detection reagent was dispensed to the wells. The plates were subsequently incubated in the dark at the RT for 1 h, and the absorbance was measured at 414 nm using a microplate reader (Varioskan LUX multimode microplate reader, Thermo Fischer Scientific, USA). Ghrelin concentrations in each plate were estimated from the standard curve. We used a linear fit because almost no curvilinear existed in the standard measures.

### Measuring blood osmolality

Blood osmolality was measured for each sample. The 250-µL plasma was extracted from each sample and the osmolality was measured using a freezing point method (Advance 3250, Advanced Instruments Inc., USA) same as that used in our previous studies (Yamada et al., 2010; Minamimoto et al., 2012; Yamada et al., 2013). The measurement error, which was evaluated as the value range for the same blood sample, was almost 2 mOsm/kgH_2_O.

### Statistical Analysis

Changes in the plasma ghrelin levels before and after eating a dry meal were analyzed using two-way analysis of variance (ANOVA) at *P* < 0.05, and plasma ghrelin (Y) was fitted using the following estimator:

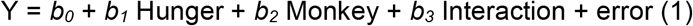

where *b*_*0*_ and the error are the intercept and residual, respectively. “Hunger” is a categorical variable to define the hunger state, differentiated by the conditions before or after food intake on a test day. “Monkey” is a categorical variable identifying the four monkeys. “Interaction” is the interaction between “Hunger” and “Monkey”. If *b*_*1*_ or *b*_*3*_ was not 0 at *P* < 0.05, we concluded that the hunger state significantly affected plasma ghrelin.

Changes in the plasma osmolality levels were analyzed using two-way ANOVA at *P* < 0.05. The plasma osmolality (Y) was fitted using the following estimator:

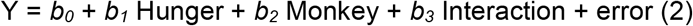

where *b*_*0*_ and the error are the intercept and residual, respectively. “Hunger” is a categorical variable to define the hunger state, differentiated by the conditions before or after food intake on a test day. “Monkey” is a categorical variable identifying the four monkeys. “Interaction” is the interaction between “Hunger” and “Monkey”. If *b*_*1*_ or *b*_*3*_ was not 0 at *P* < 0.05, then we concluded that the hunger state significantly affected the plasma osmolality.

### Influence of plasma concentration on the optical density measurements

The nonspecific effect of the plasma on the measured optical density was evaluated by adding 5% and 10% plasma to the regular concentration of the standard. All these plasma samples were obtained from a monkey on a day. We partially evaluated the degree of noise came from the plasma sample by comparing the fitted curves to these data (5% and 10%) to the standard curve without plasma (0%). We used a sample from the monkey SUN that contained relatively low ghrelin levels among the four monkeys.

The influence of the plasma concentration on the optical density measurement was evaluated using a general linear model. The optical densities (Y) were fitted using the following estimator:

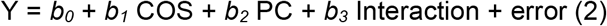

where *b*_*0*_ and the error are the intercept and residual, respectively. “COS” is a concentration of the standard. “PC” is the plasma concentration; 0%, 5%, and 10%. “Interaction” is the interaction between “COS” and “PC”. If *b*_*2*_ was not zero at *P* < 0.05, we concluded that the plasma concentration significantly affected the optical density. *b*_*2*_ should be significantly different from zero, if the plasma sample contains endogenous ghrelin. If the *b*_*3*_ was not zero at *P* < 0.05, it indicated that nonspecific effect of the plasma on the measured optical density existed in the sample.

We suspected the nonspecific effect of the plasma on the measured optical density as follows. If we assume that no endogenous ghrelin was present in the sample and that all measured optical density came from the plasma, the increase of the intercept valuess from 0% to 10% plasma indicates how much noise comes from the plasma sample. Since it is not possible to extract endogenous ghrelin from the sample, the maximum possible noise level was estimated in this way.

## Results

### Effect of dry meal intakes on blood level measurements of ghrelin and osmolality

We measured the blood levels of ghrelin and osmolality before and after consuming the regular dry meal when no water access was provided to the monkeys during an approximately two-and-a-half-hour period during and after eating the meal until the second blood collection. This controlled food access procedure mimicked a typical experimental condition to measure animal behavior in an experimental room that motivates subject performances to obtain food rewards, while its hydration status was not controlled to motivate subject performances. The amount of dry meal intake during testing was almost the same as the average amount of dry meal intake per day, indicating the animals are motivated to consume the dry meal. Two consecutive blood collections were performed on each test day with the controlled food access, which was performed a day before the test day and continued until intake after the first blood collection. Throughout the entire test period for each monkey, we did not control for water access, except for the period between the first and second blood collections. Plasma was extracted from the whole blood using a standard procedure (See Materials and Methods).

We found that the unacylated ghrelin levels tested using ELISA (See Materials and Methods) decreased on average after consuming the dry meal compared to before the intake (Fig. 1 two-way ANOVA, n = 64, d.f. = 56; Hunger, F = 5.44, *P* = 0.023; Monkey, F = 29.8, *P* < 0.001) in three out of four monkeys. Additionally, blood ghrelin levels were largely different among the monkeys (pg/ml: monkey SUN, 54–350; monkey MI, 79–380; monkey Y1, 174–1,122; monkey MO, 206– 1,123). Monkey Y1 exhibited a decrease in the value in all eight tests, and monkeys MO and MI exhibited the value decreases in six out of eight measurements. Contrastingly, monkey SUN, which exhibited the lowest blood ghrelin levels, showed the opposite effect, that is, an increase in blood ghrelin levels (Interaction, F = 3.61, *P* = 0.019) in seven of eight measures. Thus, while individual differences existed, food intake consistently decreased the blood ghrelin levels, except in one monkey.

**Fig. 1.**
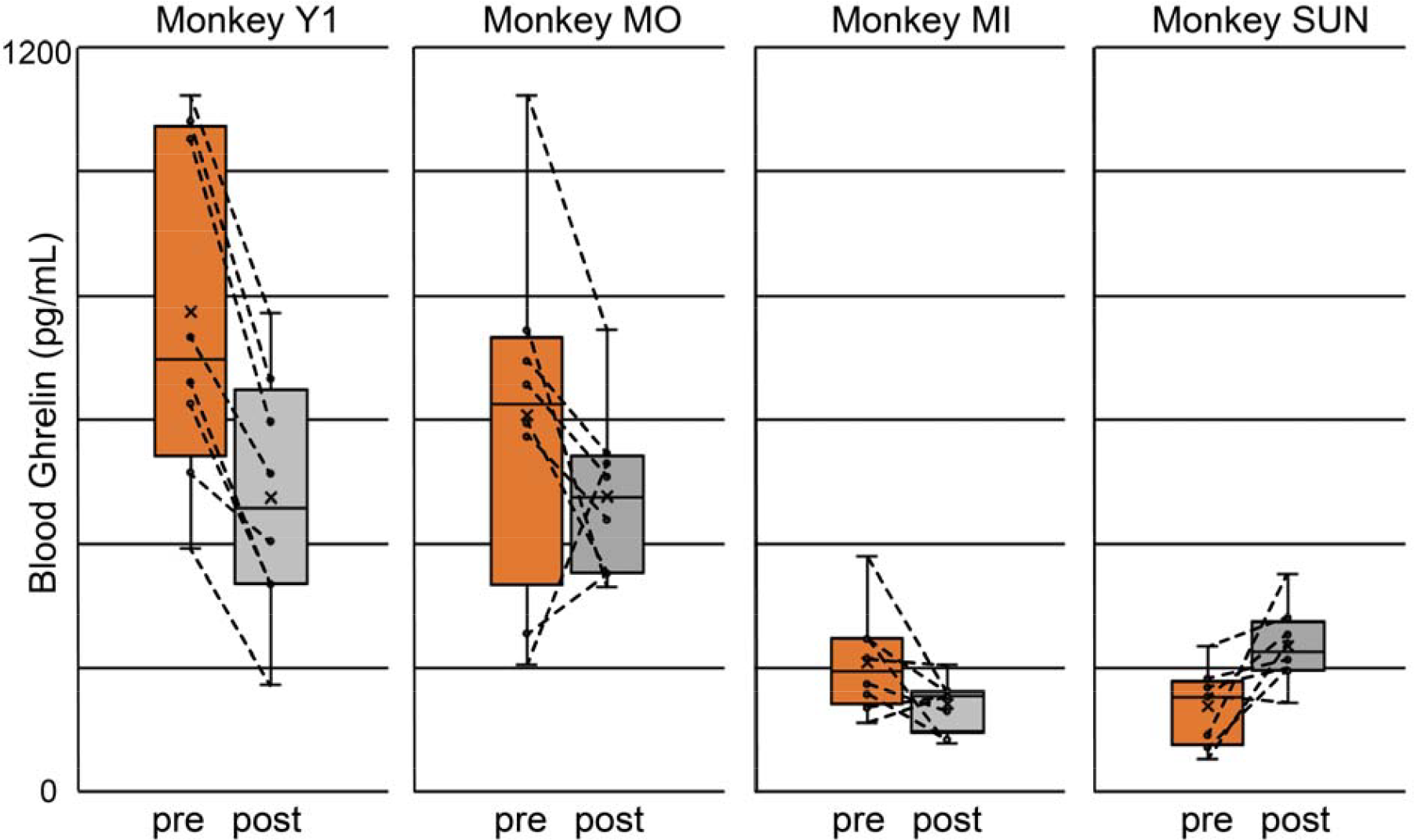
Blood ghrelin levels before and after dry meal intake. Box plot of blood ghrelin levels before (pre) and after (post) consuming regular dry meals in the four monkeys. The mean is indicated by a cross. Consecutive measurements per day were connected using a line.

To examine the effect of dry-meal intake on thirst levels, we measured blood osmolality from the same blood sample as described above. Our simple prediction for the result is that the monkeys became thirstier after eating a dry meal without fluid intake, which is a typical condition when using foods rewards to motivate monkeys in an experimental room. After dry meal intake without water, all four monkeys exhibited consistent increases in blood osmolality (Fig. 2 two-way ANOVA, n = 64, df = 56; Hunger, F = 45.7, *P* < 0.001). The effect of dry meal intake on blood osmolality clearly differed from that on the blood ghrelin levels, with no significant individual differences in osmolality levels and no significant interaction between food intake and the monkeys (Monkey, F = 2.61, *P* = 0.060; Interaction, F = 0.41, *P* = 0.746). Indeed, none of the samples in the 32 testing days (eight testing days × four monkeys) showed a decrease in osmolality. Thus, the blood osmolality level was capable of consistently describing changes in the internal state of thirst in all four animals, which is consistent with the results of our previous studies (Yamada et al., 2010; Minamimoto et al., 2012; Yamada et al., 2013).

**Fig. 2.**
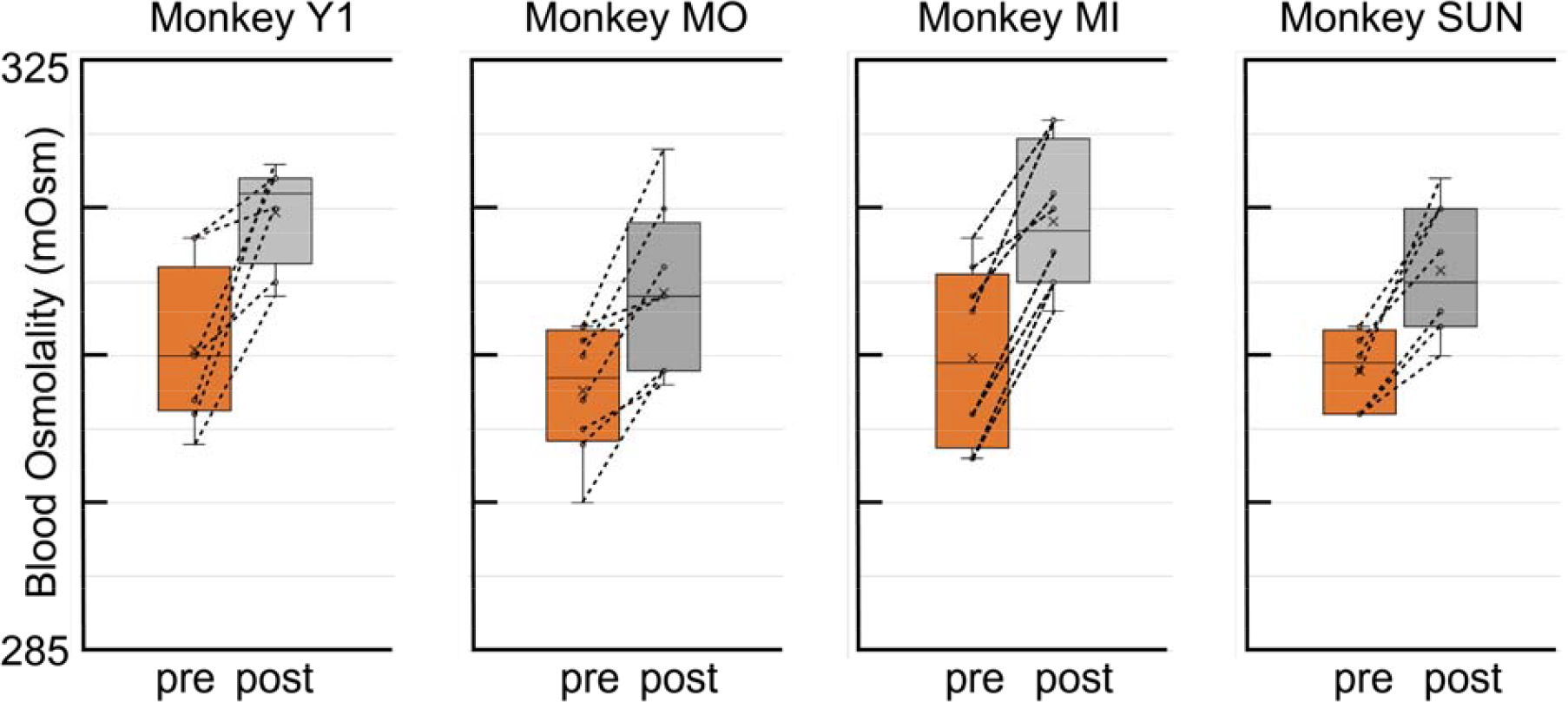
Blood osmolality levels before and after dry meal intake. Box plot of blood osmolality levels before (pre) and after (post) consuming regular dry meals in four monkeys. The mean is indicated by a cross. Consecutive measurements per day were connected using a line.

We further examined the relationship between changes in blood osmolality and ghrelin levels to directly understand whether these values were related to each other. We visualized the relationship between the changes in osmolality and ghrelin levels in each period before and after consuming the dry meal (Fig. 3), and examined whether a relationship existed between the osmolality and ghrelin levels using regression analysis (see the regression slope in Fig. 3). We found no consistent effects among the monkeys. Monkeys Y1, MO, and MI did not show consistent regression slopes, although these three individuals consistently showed a decrease in blood ghrelin levels and an increase in blood osmolality levels (see also Figs. 1 and 2). Monkey SUN, which had the opposite effect on blood ghrelin levels, showed regression coefficients similar to those in monkey MO. Thus, blood ghrelin and osmolality levels did not covary with each other either before or after consuming dry meals, indicating that these two physical measures may be able to evaluate hunger and thirst states in animals.

**Fig. 3.**
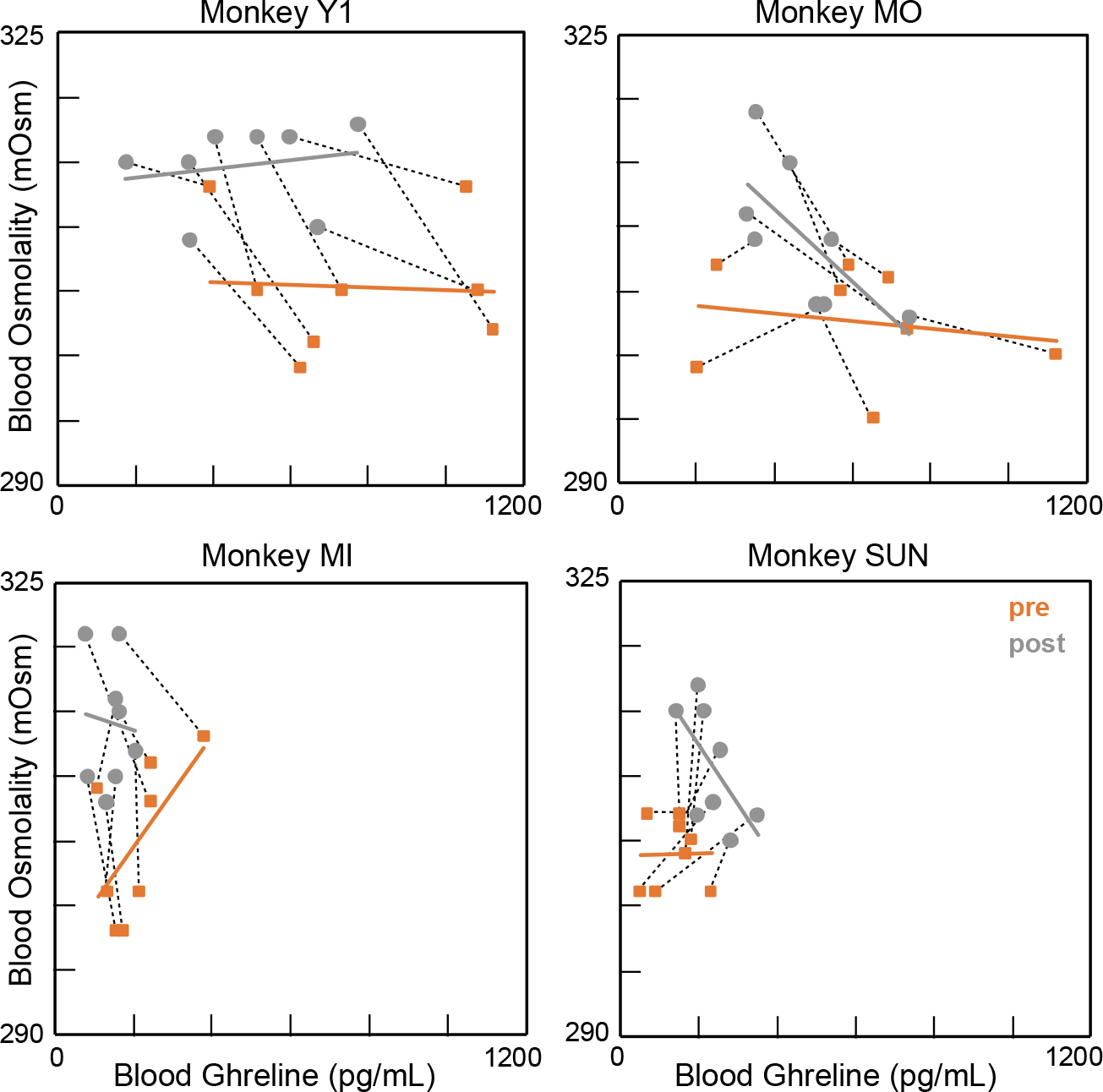
No consistent relationship was observed between the blood ghrelin and osmolality levels. Two consecutive measurements per day are represented by dotted black lines. The thick gray and orange lines indicate the regression lines for the data in the pre- and post-periods, respectively.

In summary, after dry meal intake without water, the blood ghrelin levels decreased, except in monkey SUN, and osmolality increased in all four monkeys. We did not observe a clear relationship between blood osmolality and ghrelin levels. These data indicate that the accuracy of the measures and physical changes were different across each individual.

### Assessing the influence of plasma concentration on optical density measurement

We evaluated the degree of measurement errors (or noise) included in our ghrelin measurements from the plasma, which allowed us to compare the two different physical measures of the indices for hunger and thirst with both measurement errors. For the osmolality measures we employed, we previously evaluated the measurement error, as the value range from a single sample was within almost 2 mOsm/kgH2O in our procedure. Here, we assessed the degree of error included in our optical measurement of unacylated ghrelin using ELISA. Since plasma has a nonspecific effect on optical absorption, we evaluated this noise as follows: We prepared the different concentrations of plasma samples obtained from a monkey (SUN) in one day and added them to the standard at different dilutions (0% or no endogenous plasma, 5%, and 10%). We compared these three optical measures, which contained original standard and endogenous ghrelin in 5% and 10% plasma samples. Ideally, the fitted lines changed consistently as the percentage increased. Note that removing endogenous ghrelin from plasma is impossible and that optical absorption is derived from endogenous ghrelin, plasma, and standard ghrelin.

First, all three curves were well described with a linear fit because R-squared values were > 0.99, indicating that the 5% and 10% plasma and the endogenous ghrelin elicited a consistent increase in the optical absorption (Fig. 4). The background optical absorption without ghrelin and plasma was approximately 0.141 as estimated from the fitted curves (see the intercept value of the black regression slope). These values increased to 0.192 and 0.269 in 5% and 10% ghrelin-containing buffer, respectively. These increments were 0.051 from 0% to 5% and 0.077 from 5% to 10%. This nonlinear effect was a 1.5-fold increase (0.077/0.051) and may be elicited by a 5% increase in plasma concentration from 5% to 10%. Moreover, the relative influence of the additive plasma on the optical measurements decreased as the amount of the standard ghrelin increased (almost no differences were observed at 125 pg/mL): optical densities were 0.949, 0.952, and 0.966 in 0%, 5%, and 10% plasma at the 125 pg/m, respectively. The slopes of the regression coefficients in these three linear fits became slightly shallower as the plasma concentration increased (general linear model: n = 24, d.f. = 20, concentration of standard: regression coefficient, 0.0128, t = 55.8, *P* < 0.001; plasma concentration, regression coefficient, 0.0066, t = 13.7, *P* < 0.001; interaction, regression coefficient, −0.00008, t = −4.6, *P* < 0.001). Thus, the blood ghrelin levels contained in our samples were reliably measured using 10% plasma through ELISA. In addition, some small nonspecific optical signals were derived from plasma.

**Fig. 4.**
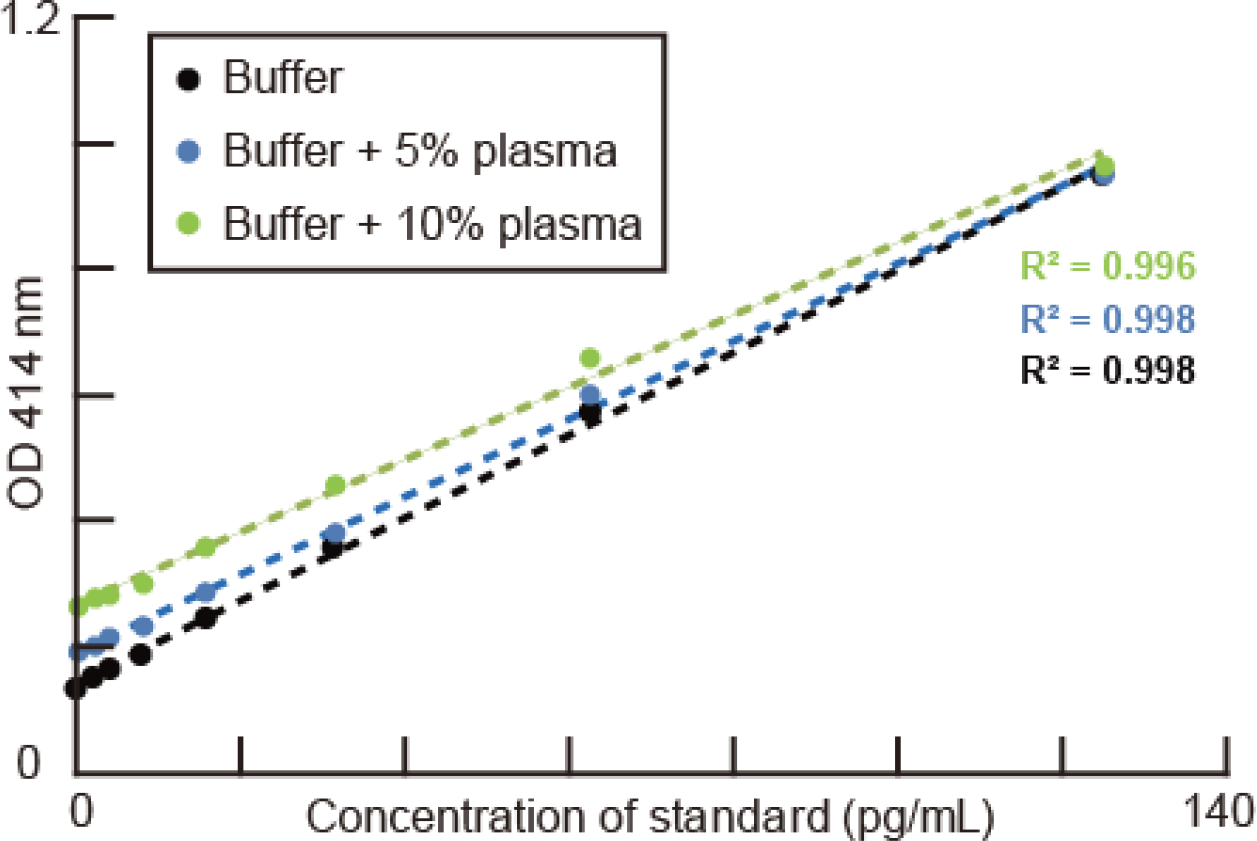
Assessing the influence of plasma concentration on measuring optical density. Plots of measured optical density (vertical axis) against standard concentrations. Each point was obtained from average of two wells, including no plasma (black), 5% plasma (blue), and 10% plasma (green). The colored dotted lines indicate the linear fit to the data. R squares are indicated.

Finally, we estimated the maximum possible noise level in the optical measures. If we assume that the no endogenous ghrelin was contained in the sample, the increase in the optical measures can serve as the noise from the plasma (i.e., maximum possible noise). This increase was estimated from the intercept differences between 0% (i.e., standard) and 10% plasma concentration, and can serve as the maximum level of noise derived from the plasma under the assumption. This increase was 0.128 (0.141 to 0.269) for 10% plasma, and thus, the value in normal plasma was 12.8 pg/mL. Since the plasma contains endogenous ghrelin, which increases optical absorption, we can only conclude that the maximal noise coming from the plasma must be smaller than this value. Thus, nonspecific optical absorption from the 10% plasma was relatively small to measure the blood ghrelin level within the range observed in these monkeys, i.e., 12.8 pg/mL noise may come from 10% plasma within the entire range of zero to 1,200 pg/mL in our measurements.

## Discussion

In this study, food consumed after 20 h of no food intake elicited a decrease in blood unacylated ghrelin levels in three out of four monkeys. Blood ghrelin levels were observed within various value ranges across individual monkeys, and each individual showed a decrease in the value, whereas one monkey showed an increase in the value. Contrastingly, the blood osmolality level perfectly reflected the change in fluid balance in all monkeys after dry meal intake; the blood osmolality level consistently increased after dry meal intake for approximately 3 h without fluid intake. Although blood osmolality and unacylated ghrelin levels can be used as indices for thirst and hunger levels, respectively, we need to consider the different characteristics of physical pressure and the peptide hormone levels as hematological indices for evaluating the two satiety states.

### Measuring ghrelin as a physical index of hunger levels

We found evidence that the decreased after dry meal intake, which reflects hunger levels (Fig. 1). This result is consistent with those of human studies showing the role of ghrelin in food intake and its changes throughout the day (Spiegel et al., 2011). In the Spiegel’s study, an increase in unacylated ghrelin levels in humans occasionally covaried with acylated ghrelin levels on average. Most previous human studies reported average changes across participants (Spiegel et al., 2011; Marchio et al., 2019). Contrastingly, our repetitive measurements for each monkey provided individual changes in ghrelin and demonstrated the reliability of blood ghrelin levels as a physical index of hunger levels.

Ghrelin acts in the brain to regulate food intake, body weight, and glucose metabolism (Tschop et al., 2000; Nakazato et al., 2001), and is a peripheral hormone released from the stomach that transmits satiety signals to the central nervous system (Sato et al., 2014). Acylated and unacylated ghrelin in humans, monkeys, and rodents are peptides consisting of 28 amino acid residues. Acylated ghrelin is the active form that modifies the third serine with the fatty acid n-octane in humans. The amino acid sequence of mammalian ghrelin is highly conserved among humans, monkeys, rats, mice, cows, pigs, and sheep. Furthermore, 10 amino acids from the N-terminus are the same for the active region (Kaiya et al., 2011), which is used in ELISA to detect endogenous ghrelin in humans and monkeys. Unacylated ghrelin is found in the stomach and blood at a certain concentration and the unacylated ghrelin does not bind to the growth hormone secretagogue receptor; however, some studies have reported that unacylated ghrelin enhances eating behavior (Toshinai et al., 2006). Thus, unacylated ghrelin may serve as a physical index of hunger levels. We note that acylated ghrelin, which is approximately 100 pg/ml in humans, was not measured in this study, and whether acylated ghrelin can be used as an index for hunger levels was unclear.

### Blood osmolality measurement as a physical index of thirst levels

We found that the fluid balance of monkeys after eating a dry meal was reflected in their blood osmolality levels. The increase in the extracellular dehydration state, as measured by the osmolality level, indicates how thirsty the monkeys were after eating the dry meal (Fig. 2). This increase in osmolality level (7 mOsmo/kgH2O) was almost equal to the osmolality increase under regular controlled water access in our previous study (8 mOsmo/kgH2O, Fig. 3 Yamada et al., 2010). In our previous study, animals worked harder to earn more water on days when they exhibited lower hydration states (higher osmolality) than when they exhibited higher hydration states (lower osmolality). This relationship between osmolality level and water consumption behavior reflects the physiology of fluid balance, as well as the increased osmolality observed in the present study.

Osmoreceptors are located at the anterior-ventral third ventricle region of the hypothalamus and detect changes in blood osmolality (Buggy et al., 1979). Neurons in the organum vasculosum of the lamina terminalis change their firing rates according to blood osmolality (Thrasher and Keil, 1987) and stimulate or suppress water and salt ingestion. In the present study, the fluid balance changed after consuming a dry meal, as the monkeys were not allowed to access water. The certified diet we fed contained water (8%), protein (21%), fat (4%), coarse ash (8%), crude fiber (3%), soluble nitrogen (53%), salt (3%), and vitamins. The monkeys ate > 100 g of this meal, which contained > 3 g of salt, and salt intake without fluid intake must stimulate and change the body fluid balance. Another important mechanism is that a reduction in blood volume stimulates fluid ingestion and suppresses diuresis through the renin–angiotensin system. (Berne and Levy, 1993). Thus, osmolality is a critical component of the hydration state, and the direct blood osmolality measurement provides a reliable index to quantify the hydration state.

### Age, sex, species, and experimental procedures

In our experiment, we used four macaque monkeys, one of which exhibited distinctive changes in blood ghrelin levels (Fig. 1 Monkey SUN). Although ghrelin plays essential roles in adiposity and metabolisms in vertebrates (Kaiya et al., 2013), the ghrelin levels were heterogeneous and different between individuals. Multiple issues may be related to this discrepancy, and we discuss the potential factors that affect blood ghrelin levels. First, monkey SUN, which exhibited an increase in blood ghrelin levels, differed from the other monkeys in age, species, and the blood collection procedure. The monkey SUN was a rhesus macaque, whereas the others were Japanese macaques, although they are closely related species, which is one possible explanation. Sex differences cannot explain our results because monkey MI is the only female among the four monkeys. Age must be the most reasonable explanation for the difference since monkey SUN was middle-aged (13 years old) compared to the other young monkeys (2-4 years old, see Materials and Methods). Ghrelin is a growth hormone and in healthy human children, total ghrelin and unacylated ghrelin levels decrease in an age-dependent manner (Wilasco et al., 2012). In aged, frail adults, lower levels of total ghrelin and its impaired response to a meal test have been observed, suggesting that ghrelin contributes to anorexia mechanisms associated with aging (Serra-Prat et al., 2009). Blood collection from monkey SUN was performed while awake, but age-related changes in ghrelin levels were most likely to explain the results observed in the monkey SUN. Additionally, a larger number of individual observations across different ages are required to elucidate this issue.

### Neuroeconomic perspectives and satiety states

Hunger and thirst are different psychological (Powloski, 1953; Zimbardo and Montgomery, 1957; Ramachandran and Pearce, 1987) and physiological phenomena (Berne and Levy, 1993); however, they are interdependent (McKiernan et al., 2009), as observed in mammals and flies (Jourjine, 2017). From an economic perspective, this interdependence may explain that food and beverages are complementary goods (Mas-Colell et al., 1995). From an food science perspective, foods and beverages contain similar materials; energy, water, salt, and other essential nutrients. Pathologically, it is associated with obesity (Martens and Westerterp-Plantenga, 2012). From a neuroscience perspective, the interdependency is explained by the facto that neural circuits for hunger and thirst interact with each other (Rolls, 1975; Jourjine, 2017; Eiselt et al., 2021; Tan et al., 2022).

In neuroeconomic studies, hunger, thirst, and their satiation, have a great impact on animal behavior (Minamimoto et al., 2012; Yamada et al., 2013; Yamada, 2017), although thirst and hunger may affect neural representations categorized as taste in the brain (Augustine et al., 2020). Physiologically, the hypothalamus plays an integral role in energy and water homeostasis by sensing and reacting to systemic signals of caloric and osmotic fluctuations (Lee, 2016). Our results observed in the body fluid balance indicated that maintaining energy and water balance is fundamental to survival; thus, a re-balance of these physiological states must occur, as the dehydration after consumption of dry meal in our testing.

For reliable estimation of animal states, both indices should be measured with similar levels of precision. We aimed to compare the measurement precision and show the index reliability with their value ranges (Fig. 4) under an assumption. We still need to improve this evaluation; however, blood ghrelin and osmolality measurements allow for a better understanding of the neural basis of decision-making as a physical index for the level of satiety. These measurements may be an easy and efficient way to evaluate the controlled access to food and fluid animals engaged in homepage.

## Acknowledgements

The authors express their appreciation to Yoshiko Yabana, Masafumi Nejime, and Shiho Nishino for their technical assistance. The authors appreciate the comments of Akira Suwa. Monkeys were provided by NBRP “Japanese Monkeys” through the National Bio Resource Project of MEXT, Japan. This study was supported by JSPS KAKENHI Grant Number JP19H05007 and Moonshot R&D JPMJMS2294 (H.Y.).

## Notes

**Conflict of interest:** The authors declare no competing interests.

### Competing Interest Statement

The authors have declared no competing interest.

